# Loss of Nrf1 rather than Nrf2 leads to inflammatory accumulation of lipids and reactive oxygen species (ROS) in human hepatoma cells, which is alleviated by 2-bromopalmitate

**DOI:** 10.1101/2021.09.29.462358

**Authors:** Rongzhen Deng, Ze Zheng, Shaofan Hu, Meng Wang, Jing Feng, Peter Mattjus, Zhengwen Zhang, Yiguo Zhang

**Author notes:** Contributed equally to this work (Rongzhen Deng, Ze Zheng).

## Abstract

Since Nrf1 and Nrf2 are essential for regulating the lipid metabolism pathways, their dysregulation has thus been shown to be critically involved in the non-controllable inflammatory transformation into cancer. Herein, we have explored the molecular mechanisms underlying their distinct regulation of lipid metabolism, by comparatively analyzing the changes in those lipid metabolism-related genes in *Nrf1α*^*–/–*^ and/or *Nrf2*^*–/–*^ cell lines relative to wild-type controls. The results revealed that loss of Nrf1α leads to lipid metabolism disorders. That is, its lipid synthesis pathway was up-regulated by the JNK-Nrf2-AP1 signaling, while its lipid decomposition pathway was down-regulated by the nuclear receptor PPAR-PGC1 signaling, thereby resulting in severe accumulation of lipids as deposited in lipid droplets. By contrast, knockout of Nrf2 gave rise to decreases in lipid synthesis and uptake capacity. These demonstrate that Nrf1 and Nrf2 contribute to significant differences in the cellular lipid metabolism profiles and relevant pathological responses. Further experimental evidence unraveled that lipid deposition in *Nrf1α*^*–/–*^ cells resulted from CD36 up-regulation by activating the PI3K-AKT-mTOR pathway, leading to abnormal activation of the inflammatory response. This was also accompanied by a series of adverse consequences, e.g., accumulation of reactive oxygen species (ROS) in *Nrf1α*^*–/–*^ cells. Interestingly, treatment of *Nrf1α*^*–/–*^ cells with 2-bromopalmitate (2BP) enabled the yield of lipid droplets to be strikingly alleviated, as accompanied by substantial abolishment of CD36 and critical inflammatory cytokines. Such *Nrf1α*^*–/–*^ led inflammatory accumulation of lipids, as well as ROS, was significantly ameliorated by 2BP. Overall, this study provides a potential strategy for cancer prevention and treatment by precision targeting of Nrf1, Nrf2 alone or both.

## 1. Introduction

Lipids are essential for energy storage, assembles into macromolecular structures as biomembranes, and function in cell growth, proliferation, differentiation, and movement. Many types of lipids act as biological signaling molecules and thus exert diverse functionality in energy metabolism and cell homeostasis [1-3]. However, abnormal lipid metabolism is connected to a variety of diseases. For instance, obesity caused by over-nutrition, nonalcoholic steatohepatitis (NASH), and type 2 diabetes, are all related to the imbalance in the lipid metabolism status [4]. Importantly, the development of hepatocellular carcinoma (HCC) is also linked to lipid disorders, such as NASH [5]. This is due to the fact that most cancer cells had been shown to have a higher rate of *de novo* lipid synthesis, leading to an obvious increase in newly synthesized fatty acids, glycerophospholipids and cholesterol. These serve as raw materials to meet the increasing requirements of cancer cells for their malignant growth and proliferation. In addition, the up-regulated catabolism of lipids by β-oxidation in the mitochondria can also provide additional energy, as required by those cancer cells [6].

In almost all living organisms, lipids are stored in the form of triglycerides and cholesterol-esters in lipid droplets (LDs). This hydrophobic organelle with a surrounding phospholipid monolayer membrane, can mobilize fatty acids during specific life processes [7]. Since the liver is the main metabolic organ of lipids in vertebrates, lipids play a vital role in the physiological homeostasis of the liver and related pathological processes of many liver diseases. The liver-specific mouse knockout of *Nrf1* (also called *Nfe2l1*, encoding a ‘living fossil-like’ conserved membrane-bound transcription factor of the cap’n’collar (CNC) basic-region leucine zipper (bZIP) family [8]) leads to spontaneous development of NASH and its ensuing malignant transformation into hepatoma [9]. Such hepatocyte-specific loss of *Nrf1* caused up-regulation of the cytochrome p450 family *CYP4A* and *CYP1A1* genes, hepatic microsomal lipid peroxidation to be substantially increased, and then fatty acids and cystine/cysteine contents to be aberrantly accumulated, as consequently followed by severe steatosis, apoptosis, necrosis, inflammation, and fibrosis, eventually worsening and developing into liver cancer [10, 11]. During this pathogenesis, significant activation of two lipid metabolism-related genes, *Lipin1* and *PGC-1β* was observed in the murine liver-specific *Nrf1*^*–/–*^ cells, implying that Nrf1 negatively regulates these two genes by directly binding to the putative antioxidant response elements (AREs) located in their promoter regions [12].

Nrf2 (also encoded by *Nfe2l2*, a soluble transcription factor of the CNC-bZIP family) has been shown to regulate lipid metabolism by mediating the expression of several lipid metabolism-related genes, such as those encoding ACLY, ACC1 and ELOVL6 [13, 14]. It was found by Huang et al [15] that *Nrf2*^*–/–*^ mice manifested with a loss of their liver weight at six months of age, together with a marked decrease of the triacylglycerol contents in the liver. These changes in the liver of *Nrf2*^*–/–*^ mice had led to the up-regulation of lipogenic genes including *PPARγ, FASN, SCD1* and *SREBP1*, albeit they were proved to serve direct target genes of Nrf2 [15]. Adipose-specific ablation of *Nrf2* in mice can also cause a transient delay of high-fat diet-induced obesity by altering glucose, lipid, and energy metabolism in such mice [16]. Taken together, these findings demonstrate that Nrf2 acts as an important regulator of lipid metabolism. However, how Nrf1 and Nrf2 differentially govern (and reciprocally regulate) the lipid metabolism process, in particular liver cancer cells, is still poorly understood to date.

In this study, we focus on the lipid synthesis and catabolism, as well as the mechanism related to lipid deposition, by conducting various experiments to determine the yields of lipid droplets in *Nrf1α*^*–/–*^ cells (in which the full-length Nrf1 of 742 aa was deleted by its gene editing) and its changes in the lipid metabolism gene expression, as compared to the data obtained from in *Nrf2*^*–/–*^ cells. The results unveiled that Nrf1 and Nrf2 have performed their respective regulatory functions in the lipid metabolism of hepatoma cells through distinct molecular mechanisms. Further experiments also revealed that both CNC-bZIP factors are differentially involved in putative cytoprotection against inflammatory response to abnormal lipid metabolism caused by *Nrf1α*^*–/–*^ and/or *Nrf2*^*–/–*^. Interestingly, lipid deposition and related inflammation responses are significantly reduced by a palmitic acid analogue, 2-bromopalmitate (2BP). Collectively, this study provides a novel potential strategy for the prevention and treatment of NASH and hepatoma by precision targeting of Nrf1, Nrf2 alone or both.

## 2. Materials and Methods

### 2.1 Cell lines and culture

The human HepG2 cells (i.e., *Nrf1/2*^+/+^) were obtained from the American Type Culture Collection (ATCC, Manassas, VA, USA). *Nrf1α*^*–/–*^ and *Nrf2*^*–/–*^ cell lines were established on the base of HepG2 cells in our laboratory by Qiu et al [17]. Besides, Nrf1α-restored cell line was created by employing lentivirus technology to allow for stable expression of Nrf1 αproteins and also identified to be true. The fidelity was further confirmed by its authentication profiling and STR (short tandem repeat) typing map (Shanghai Biowing Applied Biotechnology Co., Ltd). All these cell lines were cultured at 37°C with 5% CO2 in Dulbecco’s modified Eagle’s medium (DMEM) containing 25 mmol/L glucose, 10% (*v*/*v*) fetal bovine serum (FBS), 100 units/mL penicillin-streptomycin, before subsequent experiments were performed.

### 2.2 Real-time quantitative PCR analysis

The total RNAs were extracted from experimental cells when they reached 80% confluency using an RNA extraction kit (TIANGEN, Beijing, China) and then reversely transcribed into single-stranded cDNAs by a Reverse-Transcription Kit (Promega, USA). In parallel experiments, the cell lines that were subjected to different treatments with specific inhibitors or siRNAs (to interfere with its cognate target gene expression) were subjected to similar RNA extraction procedure. The mRNA expression levels of each target gene were determined by real-time quantitative PCR (i.e., RT-qPCR) with each of those indicated pairs of their forward and reverse primers (listed in Table 1).

**Table 1.**
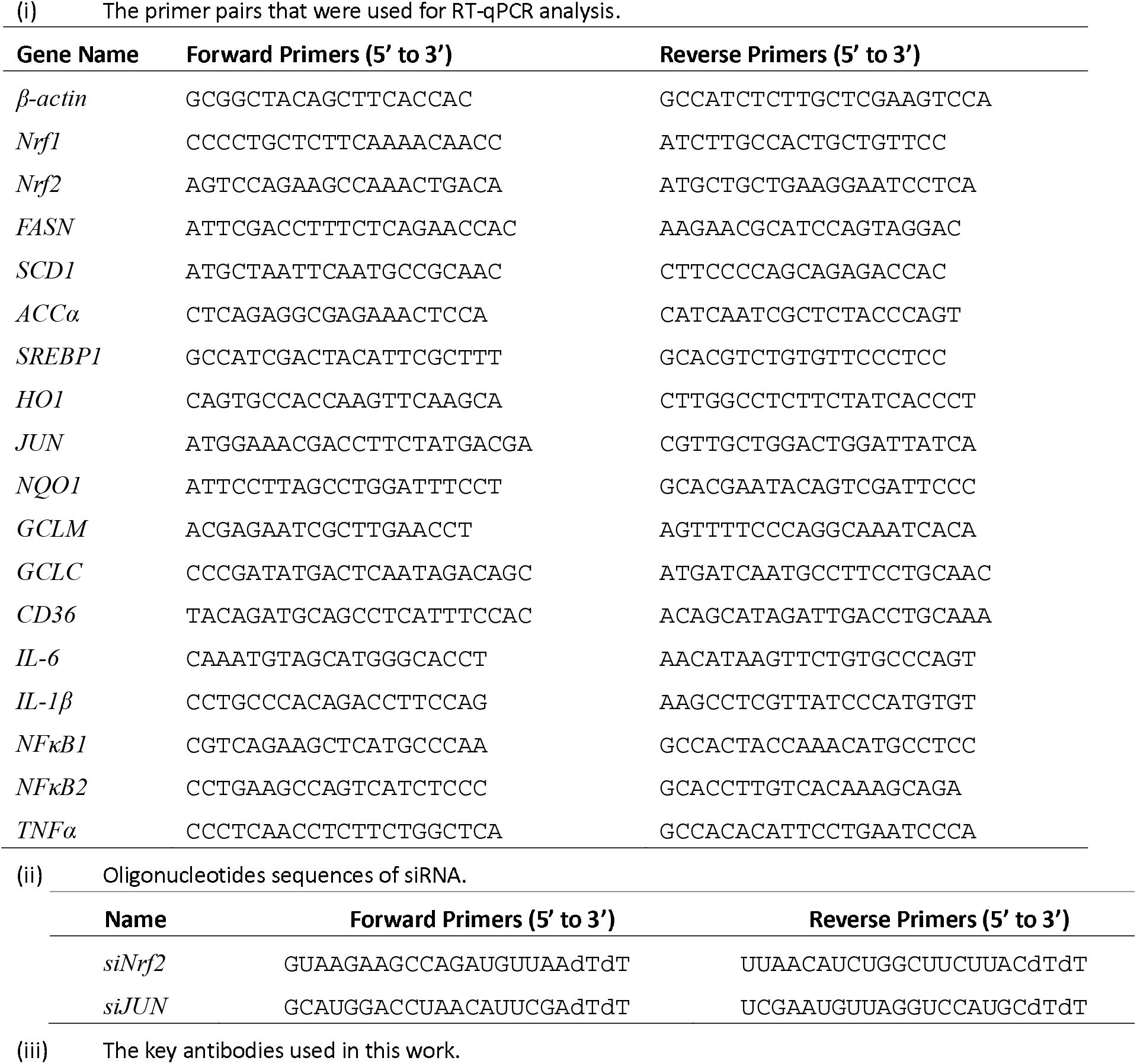
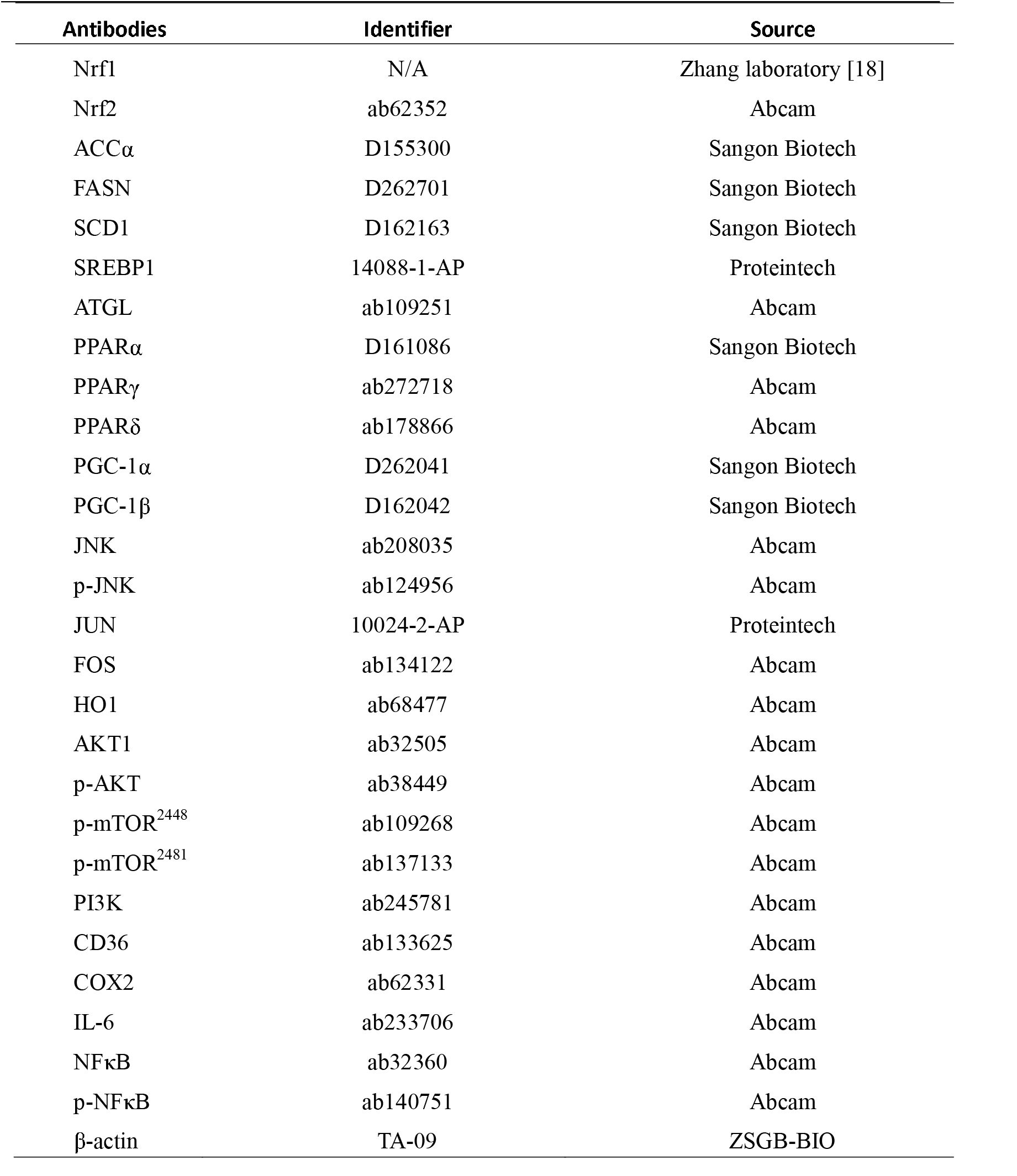
All oligonucleotides and antibodies that were used in this study.

### 2.3 Small interfering RNAs and kinase-specific inhibitors

To determine the effect of a given gene on the downstream signaling, the JNK function inhibitor SP600125 (Selleck, Shanghai, China), the PI3K inhibitor NVP-BKM120 (Cayman Chemical, USA), and the mTOR inhibitor RAPA (rapamycin, Sigma, USA) were used here. Two pairs of small-interfering RNAs (siRNAs, with their oligonucleotide sequences listed in Table 1) were designed for specifically targeting Nrf2 or JUN (i.e., siNrf2 or siJUN), in order to interfere with their mRNA expression levels (Tsingke, Beijing, China).

### 2.4 Western blotting analysis of key proteins

All experimental cells were subjected to extraction of total proteins in a lysis buffer containing a protease inhibitor cOmplete Tablets EASYpack or phosphatase inhibitor PhosSTOP EASYpack (each tablet per 15 mL buffer, Roche, Basel, Switzerland). The proteins in cell lysates were denatured at 100°C for 10 min, followed by sonication, and then diluted in 3×loading buffer (187.5 mmol/L Tris-HCl, pH 6.8, 6% SDS, 30% Glycerol, 150 mmol/L DTT, and 0.3% Bromphenol Blue) at 100°C for 5 min. Thereafter, equal amounts of cell lysates were separated by SDS-PAGE gels (8-12% polyacrylamide) and the specific proteins visualized by Western blotting with antibodies listed in Table 1.

### 2.5 Determination of intracellular triglyceride contents

All the relevant experimental cells were subjected to determination of the triglyceride (TG) contents, as conducted according to the manufacture’s instruction using a TG-enzymatic assay kit (APPLYGEN, Beijing, China). After discarding the medium and rinsing the cells with PBS for three times, 170 μL of a working solution (prepared by mixing reagent R1 with reagent R2 at the ratio of 4:1 on ice) was added into every well of 96-wells plate. And then this plate was incubated at room temperature for 15 min, before quantitative absorbance was measured at 570 nm by using spectrophotometer (Thermo Scientific, Shanghai, China).

### 2.6 Oil Red O-staining of lipid droplets in cells

After the cells reached 70% of confluency, they were subjected to different treatments with different concentrations of oleic acid (OA, Solarbio, Beijing, China) or 2-bromopalmitate (2BP, Sigma, USA). Then, the cells were fixed in a tissue fixative buffer containing 4% paraformaldehyde (Boster Biological Technology, Wuhan, China) for 30 min, followed by lipid droplet staining for 30 min in a solution containing 3 g/L of oil red O (Sangon Biotech, Shanghai, China). Finally, the stained cells were rinsed three times with 60% of isopropyl alcohol (Kelong, Chengdu, China), and the red lipid droplets were visualized and photographed by microscopically imaging analysis.

### 2.7 Assays of ARE-driven luciferase reporter gene activity

Equal numbers (2.0 × 10^5^) of experimental cells (*Nrf1/2*^+/+^) were seeded into each well of the 12-well plates. After reaching 80% confluence, the cells were co-transfected using a Lipofectamine 3000 mixture with each of lipid catabolism related *ARE*-driven luciferase reporters (constructed by inserting each of the *ARE*-battery sequences of lipid catabolism related gene into the pGL3-Promoter vector) or non-*ARE* plasmids (as a background control), together with an expression construct for Nrf1 or empty pcDNA3.1 vector. In the same time, the Renilla expression by pRL-TK plasmid served as an internal control for transfection efficiency. Approximately 24 hours after transfection, the reporter activity was measured by using the dual-luciferase reporter assay kit (Promega, USA). The resulting data were calculated as fold changes (mean ± S.D., n = 9), relative to the basal activity (at a given value of 1.0) obtained from the transfection of cells with an empty pcDNA3.1 and each of ARE-driven reporter genes. These ARE-core sequences are shown in Table 2.

**Table 2.**
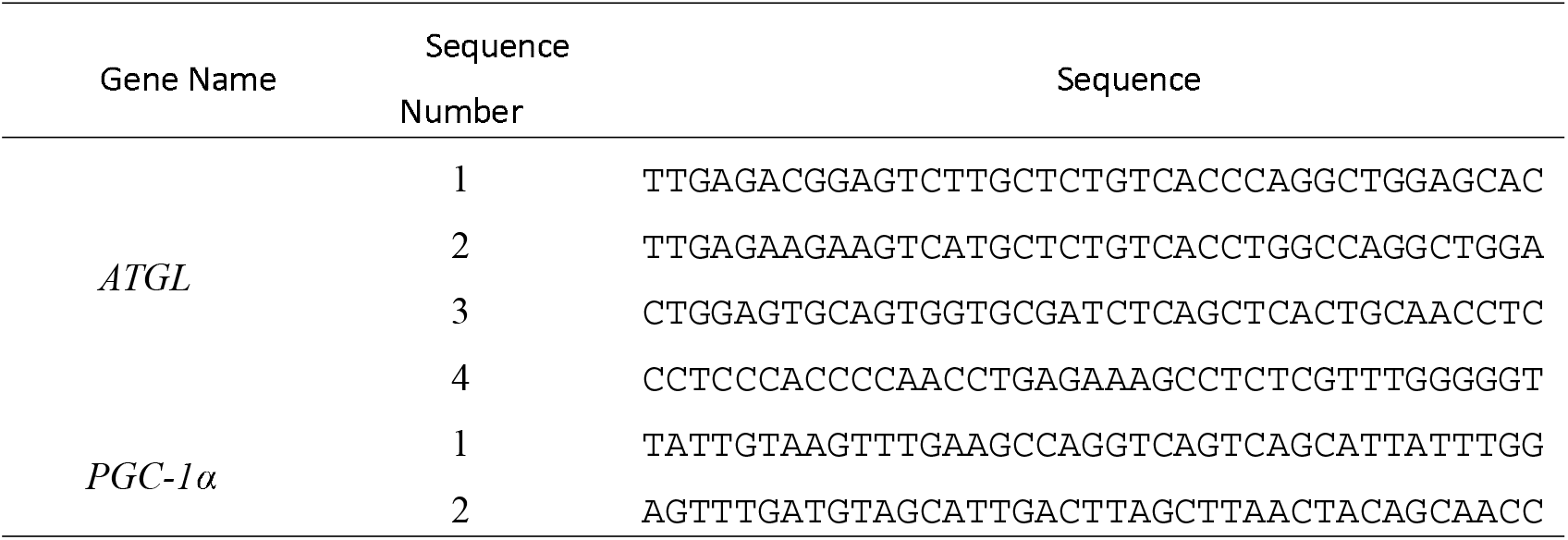
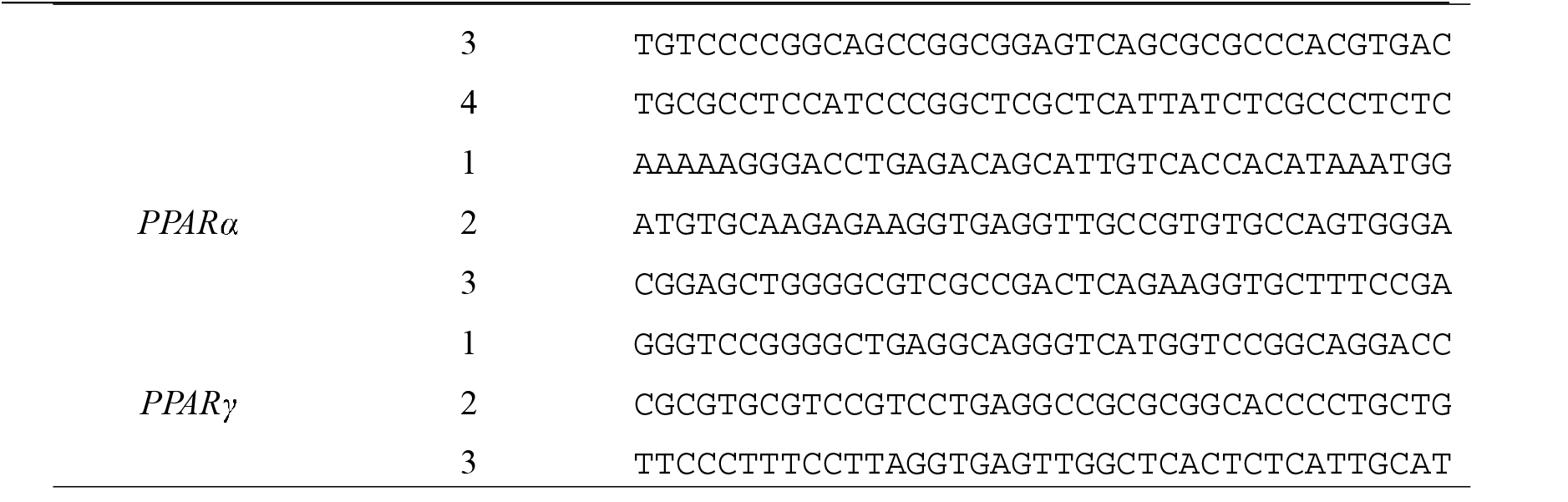
The ARE-drive reporter sequences from the lipid catabolism related gene promoters. Sequence

### 2.8 ChIP-sequencing analysis

The CHIP-sequencing data for Nrf1 (called Nfe2l1) binding to the promoter region of interested target genes in HepG2 cells obtained from the Encode database (https://www.encodeproject.org) with the project number ENCSR543SBE), were analyzed herein.

### 2.9 Analysis of cellular reactive oxygen species (ROS) by flow cytometry

Equal numbers (3.0 × 10^5^) of experimental cells were seeded into each well of the 6-well plates. At 70% confluency, they were transferred to a fresh medium with or without 2BP for 24 h and then prepared according to the manufacture’s instruction for the ROS assay kit (Beyotime, Shanghai, China). The cells were collected and suspended in diluted DCFH-DA (which was diluted with serum-free medium at 1:1000 to achieve a final concentration of 10 μM), before being incubated for 20 min at 37°C in a cell incubator. Mix upside down every 3-5 min so that the probe is allowed for a full contact with the cells, before they were then washed three times with a serum-free medium. Thereafter, the intracellular ROS levels were determined by flow cytometry (at Ex/Em = 488/525nm). The resulting data were analyzed by the FlowJo 7.6.1 software, as described previously [19]. Similar experiments were obtained by using a Mito-Tracker Red CMXRos ROS test kit (Beyotime, Shanghai, China).

### 2.10 Statistical analysis

Significant differences for the obtained data were statistically determined using the Student’s *t*-test or Multiple Analysis of Variations (MANOVA). The data are shown herein as a fold change (mean ± S.D.), each of which represents at least three independent experiments that were each performed in triplicates.

## 3. Results

### 3.1. Knockout of Nrf1α in hepatoma cells leads to lipid deposition while knockout of Nrf2 leads to lipid reduction

A striking feature in the pathology of NASH is lipid deposition, which is also a hall-mark of liver cancer development in murine *Nrf1*^*–/–*^-disordered lipid metabolism [10, 11]. Herein, wild-type HepG2 cells (*Nrf1/2*^+/+^) and two derived cell lines (*Nrf1α*^*–/–*^ and *Nrf2*^*–/–*^) were employed in the following experiments to investigate distinct functionality of Nrf1 and Nrf2 in human intracellular lipid metabolism. These cell lines were further validated by Western blotting and real-time qPCR analyses of Nrf1 and Nrf2 (Fig. 1, A & B). The results were fully consistent with our previous data [17, 19] when comparing the wild-type *Nrf1/2*^+/+^ controls to *Nrf1α*^*–/–*^ cells. Such *Nrf1α*^*–/–*^ cells exhibited no expression of Nrf1α-derived proteins, but other shorter isoforms of Nrf1ß and Nrf1γ were unaltered (Fig.1A, *upper panel*). By contrast, Nrf2 protein amounts (Fig.1A, *middle panel*) and its basal mRNA levels (Fig. 2B) were significantly increased in *Nrf1α*^*–/–*^ cells. Nrf1α-derived proteins were slightly enhanced in *Nrf2*^*–/–*^ cells lacking its two major proteins (Fig. 1A, *right lane*), but this was accompanied by a modest decrease in the basal mRNA expression of Nrf1 (Fig. 1, A & B). These indicate putative inter-regulation of Nrf1α and Nrf2 that could be executed at distinct levels.

**Figure 1.**
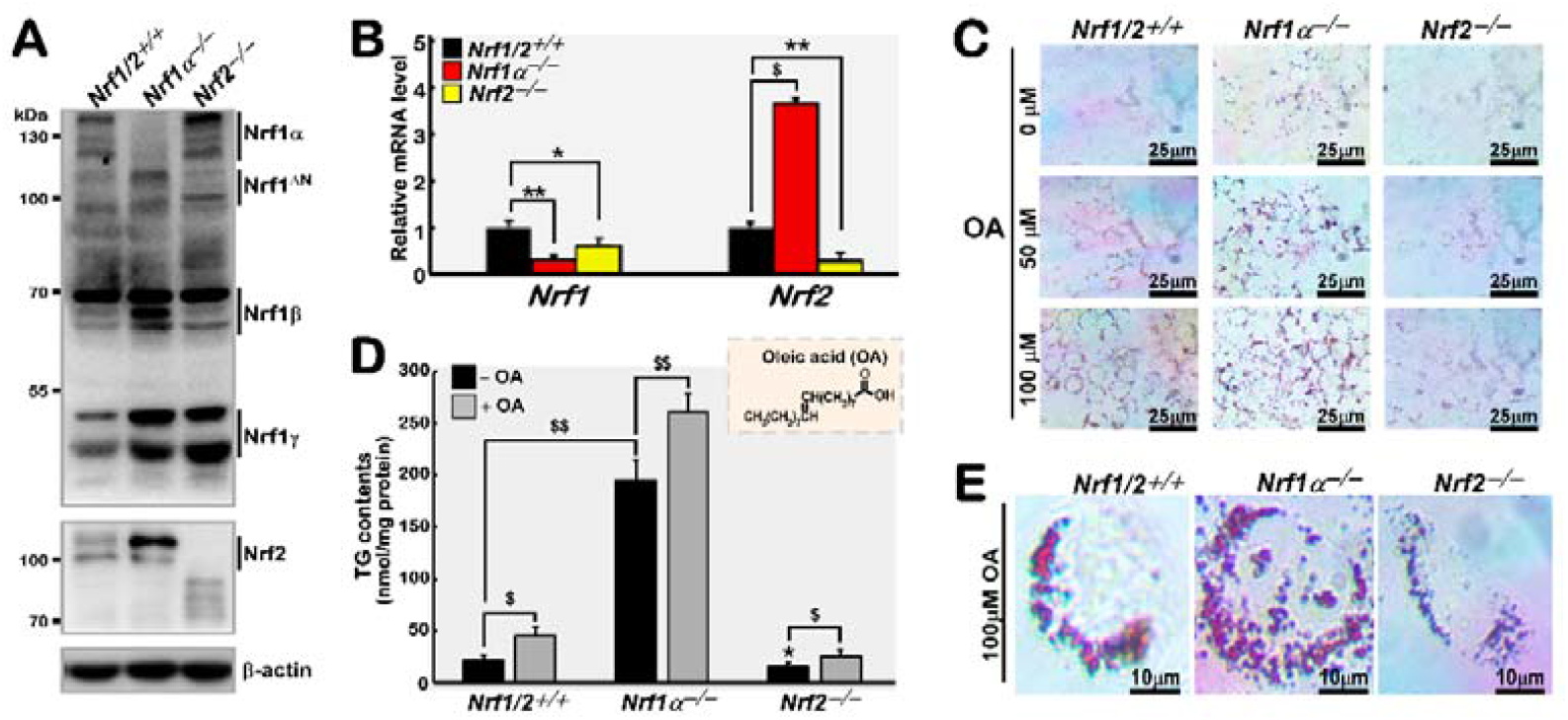
Significant differences between Nrf1 and Nrf2 in lipid metabolism are observed in their genotypic knockout cell lines. (A) Nrf1 and Nrf2 protein levels in three different genotypic cell lines *Nrf1/2*^+/+^, *Nrf1α*^*–/–*^and *Nrf2*^*–/–*^ were determined by Western blotting using specific antibodies. β-actin was used as a loading control. (B) mRNA expression levels of both Nrf1 and Nrf2 were examined by real-time qPCR analysis of *Nrf1/2*^*+/+*^, *Nrf1α*^*–/–*^ and *Nrf2*^*–/–*^ cell lines. The results were then calculated as fold changes (mean ± S.D) with significant increases ($, P < 0.01) and significant de-creases (*, *P* < 0.05; **, *P* < 0.01). (C) The triglyceride (TG) contents were measured after *Nrf1/2*^+/+^, *Nrf1α*^*–/–*^ and *Nrf2*^*–/–*^ cell lines were pretreated with 100 μmol/L oleic acid (OA). The results were then calculated as a concentration of nmol per mg of protein, with significant increases ($, *P* < 0.01; $$, *P* < 0.001) or significant decreases (*, *P* < 0.05). (D) The same cell lines were further subjected to pretreatment with distinct concentrations of oleic acid (OA), followed by the oil red O staining, and photographed using a microscope (scale bar = 25 μm), with higher resolution imaging of the same cells that had been treated as described above (E, scale bar = 10 μm). The statistical significances were determined by using the Student’s *t*-test or MANOVA.

**Figure 2.**
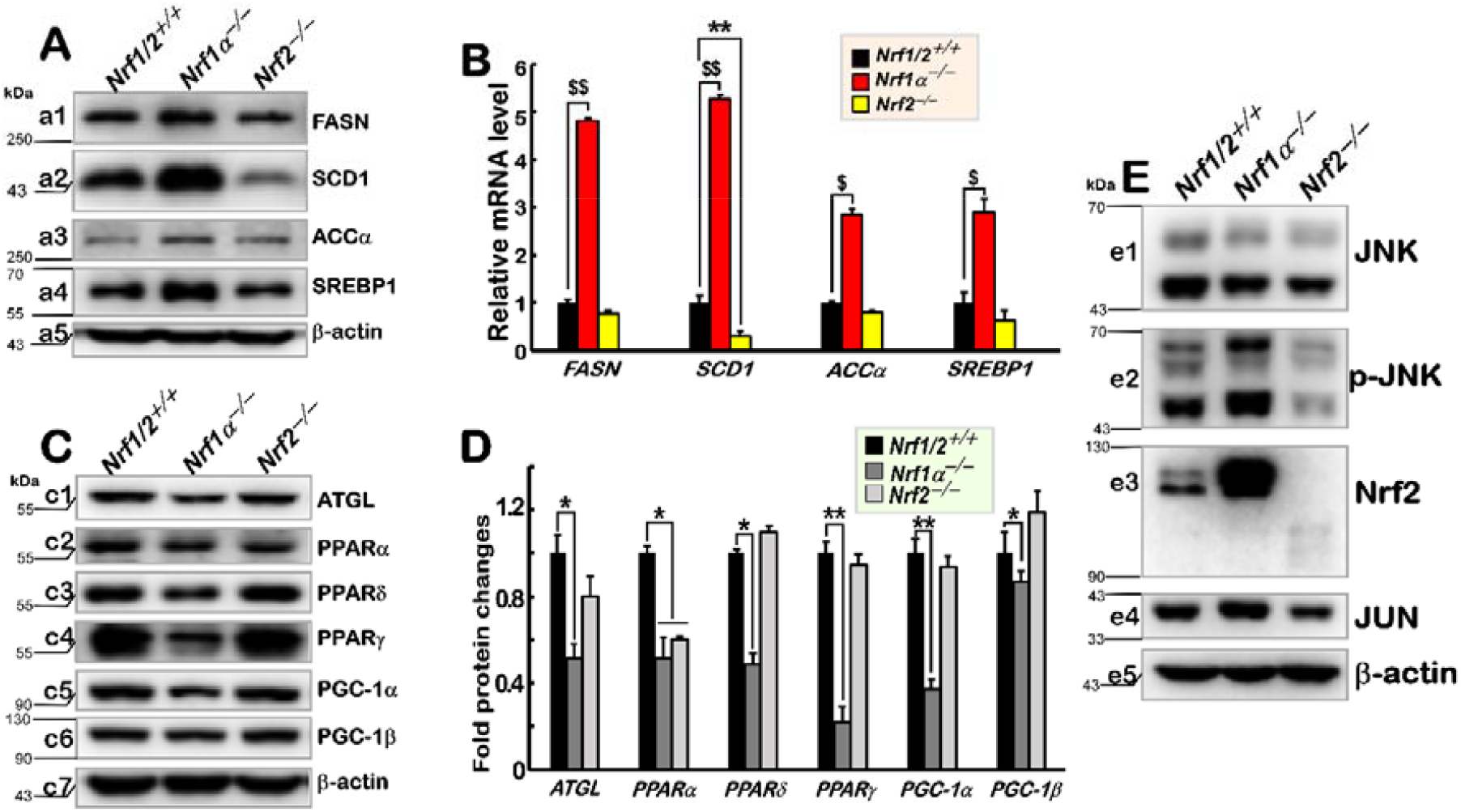
Different expression levels of lipid metabolism-related genes mediated by Nrf1 and Nrf2 in distinct genotypic cell lines. (A) Different protein levels of key genes involved in the de novo biosynthesis of lipids in *Nrf1/2*^+/+^, *Nrf1α*^*–/–*^and *Nrf2*^*–/–*^ cell lines were determined by Western blotting. β-actin was used as a loading control. (B) Different mRNA expression levels of the indicated genes were determined by real-time qPCR. The results were calculated as fold changes (mean ± S.D) with significant increases ($, *P* < 0.01; $$, *P* < 0.001) or significant decreases (**, *P* < 0.01). (C) The protein expression abundances of those key enzymes and transcription factors involved in the lipid catabolism were evaluated by Western blotting. (D) The intensity of all indicated immunoblots was calculated at fold changes (mean ± S.D) and shown graphically with significant decreases (*, *P* < 0.05; **, *P* < 0.01). (E) Basal protein expression levels of JNK, p-JNK, Nrf2 and JUN were also examined in *Nrf1/2*^+/+^, *Nrf1α*^*–/–*^and *Nrf2*^*–/–*^ cell lines. The statistical significances were determined by using MANOVA.

After 24-h treatment of distinct cell lines with oleic acid (OA) at different concentrations (0, 50, 100 μmol/L), the lipid droplets were examined by oil red O staining. The data is shown in Fig. 1C. When compared with wild-type *Nrf1/2*^+/+^ controls, much more and larger lipid droplets were found in *Nrf1α*^*–/–*^ cells, whereas small lipid droplets were in *Nrf2*^*–/–*^ cells. There was a significant difference in the visualization of lipid droplets after OA treatment of these three cell lines with incrementing concentrations of OA from 50 to 100 μmol/L (Fig. 1C). Such an OA-increased trend of lipid droplets was seen in *Nrf1α*^*–/–*^ cells that were significantly higher than that of *Nrf1/2*^+/+^ controls, while *Nrf2*^*–/–*^ cells also manifested a reduced visualization of lipid droplets. These unraveled that loss of *Nrf1α* led to an aberrant accumulation of lipids, whereas similar lipid deposition appeared to be alleviated by inactivation of Nrf2. This finding was also further evidenced by an intracellular triglyceride (TG) enzymatic detection assay of those same cell lines that had been treated with 100 μmol/L OA (Fig. 1D). Their distinct TG contents (in lipid droplets) were also visualized by oil red O staining, as shown in Fig. 1E. These distinct results demonstrated that loss of Nrf1α enables the lipid deposition to be augmented in *Nrf1α*^*–/–*^ hepatoma cells, but a similar lipid-accumulating phenomenon was not observed in *Nrf2*^*–/–*^ cells. The latter further implies that its lipid synthesis and/or uptake may have been suppressed by *Nrf2*-specific knockout of its transactivation domain (i.e., *Nrf2*^*––*^*Δ*^*TA*^) rather than its DNA-binding domain as described previously [20]

### 3.2 Loss of Nrf1α or Nrf2 causes different effects on the expression of genes responsible for lipid metabolism

It has been shown that genes governing the de novo lipid synthesis were expressed in cancer cells to a considerably greater extent than their equivalents in normal cells [21]. Therefore, a few of key rate-limiting enzymes (e.g., FASN, ACCα, SCD1) of de novo lipid synthesis and regulated transcriptional factor SREBP1 (sterol regulatory element-binding protein 1) were examined next. As anticipated, our results revealed that the loss of *Nrf1α*^*–/–*^ caused a substantial increase in the protein expression of FASN (fatty acid synthase), ACCα (acetyl-CoA carboxylase alpha), SCD1 (stearoyl-Coenzyme A desaturase 1) and SREBP1 when compared to their counterparts in wild-type *Nrf1/2*^+/+^ cells (Fig. 2A). Similar augmented expression levels of mRNA were also observed in *Nrf1α*^*–/–*^ cells when analyzed using real-time qPCR (Fig. 2B). Conversely, the lentivirus-mediated recovery of Nrf1α in *Nrf1α*^*–/–*^ cells (i.e., Nrf1α-restored cell line, as illustrated in Figs. S1 & S2A) enabled to cause a substantial decrease in the protein expression of FASN, ACCα, SCD1 and SREBP1 when compared to *Nrf1α*^*–/–*^ cells (Fig. S2B). These indicate that aberrant expression of lipid synthesis-related genes is caused by the absence of *Nrf1α*. By striking contrast, *Nrf2*^*–/–*^ cells gave rise to down-regulated expression levels of the same genes to greater or fewer degrees (Fig. 2, A & B).

Further examinations of *Nrf1α*^*–/–*^ cells revealed a considerably lower expression level of adipose triglyceride lipase (ATGL, also called PNPLA2, which is the catalytic rate-limiting enzyme in the first step of TG hydrolysis during oxidation catabolic mobilization [22], Fig. 2C & D). A small decrease in the expression of ATGL was also determined in *Nrf2*^*–/–*^ cells as compared to wild-type cells (Fig. 2C & D). Further experimental analysis of the nuclear receptor-types (i.e., PPARα, γ, δ) of transcription factors governing oxidative degradation of fatty acids [23] and their interacting transcriptional co-activators (i.e., PGC-1α, β) [24] was carried out. The results showed that all the examined proliferator-activated receptors (PPARα, γ, δ) and PPARγ co-activators (PGC-1α, -1β) were significantly down-regulated in *Nrf1α*^*–/–*^ cells (retaining a hyper-active Nrf2), and thus were roughly unaffected in *Nrf2*^*–/–*^ cells, but except for down-expressed PPARα only (Fig. 2, C & D). Conversely, ATGL, PGC-1α and PPARα were significantly up-regulated in Nrf1α-restored cells, as compared with *Nrf1α*^*–/–*^ cells (Fig. S2C). Such differential expression patterns of PPARα, γ and δ regulated by Nrf1 and/or Nrf2 implies a selectivity to mediate distinct downstream genes, and thus it is inferable that PPARγ and PPARδ are the main downstream of Nrf1α, whereas PPARα is not. Conversely, upon loss of Nrf1α, the former two factors will be down-regulated, while the latter one has no significant changes. Taken altogether, these collective data demonstrate that abnormal deposition of lipids in *Nrf1α*^*–/–*^ cells result from up-regulation of lipid anabolism, along with substantial down-regulation of lipid catabolism-controlled genes, but the opposite effects seem to be obtained in *Nrf2*^*–/–*^ cells.

Next, the differential expression profiles of lipid synthesis- and catabolism-related genes between *Nrf1α-* and *Nrf2*-deficient cell lines were further explored. The results showed a markedly increased presence of phosphorylated JNK and its cognate target JUN in *Nrf1α*^*–/–*^ cells (with a hyper-active Nrf2) (Fig. 2E, *e2* to *e4*), although the total protein level of JUK was slightly down-regulated (Fig. 2E, *e1*) when compared to wild-type controls. However, Nrf1α-restored cells only manifested a modest decreased expression level of JUN (Fig. S2C). By contrast, all those examined protein levels were strikingly diminished upon inactivation of *Nrf2*^*–/–*^ (Fig. 2E). Such similarly-altered expression trends of Nrf2, p-JUNK and JUN seem to presage a comparable pattern to those of the lipogenic genes, thereby providing a new clue for the further study of their relevant gene regulation.

### 3.3 The JNK-Nrf2-AP1 signaling is required for up-regulation of lipogenic and HO-1 genes in Nrf1α-deficient cells

As shown in Fig. 3A (*a1* to *a8*), treatment of *Nrf1α*^*–/–*^ cells with a JNK-specific inhibitor SP600125 (to block the activity of phosphorylated JNK) caused substantial down-regulation of all seven other examined proteins, including Nrf2, JUN (a direct substrate of JNK), HO-1, FASN, ACCα, SCD1 and SREBP1. Notably, the lipid synthesis-related ACCα and SCD1 displayed a profound inhibitory trend similar to those of JUN and Nrf2, but SREBP1 appeared to be only partially inhibited by SP600125. This implies that the JUK signaling to Nrf2 and/or AP1 is involved in the aberrant lipogenesis of *Nrf1α*^*–/–*^ cells. The treated cells still retained a remnant fraction of JUN and Nrf2, but their direct downstream HO-1 expression was almost completely abolished by SP600125 (Fig. 3A, *cf. a4* with *a2* and *a3*), implicating an integrative effect of both factors on HO-1.

**Figure 3.**
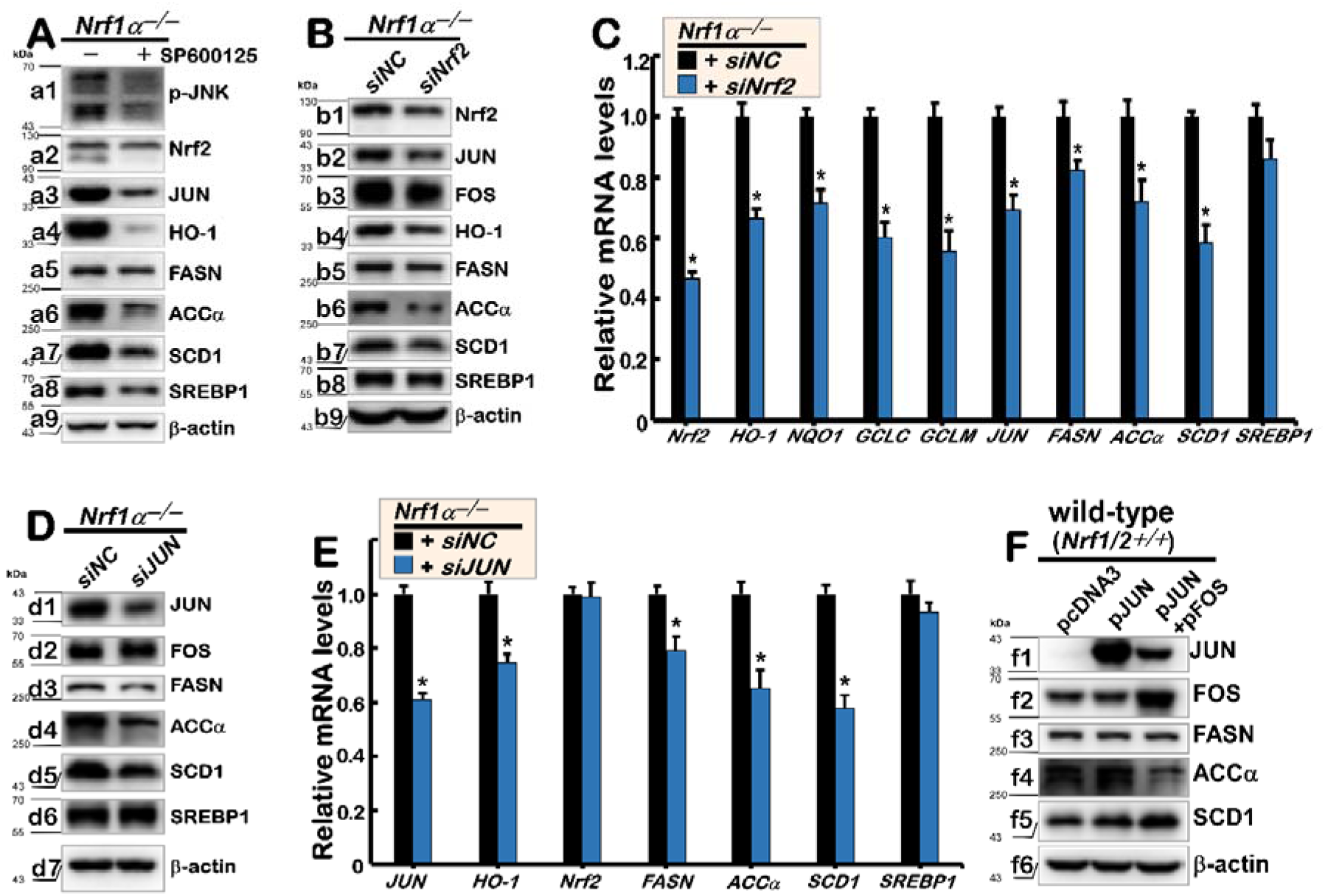
The JNK-Nrf2-AP1 signaling pathway is required for increased expression of lipid synthesis genes in *Nrf1α*^*–/–*^ cells. (A) *Nrf1α*^*–/–*^ cells were treated with or without the JNK-specific inhibitor SP600125 (20 μmol/L) for 24 h and then subjected to Western blot analysis of the JNK-Nrf2-AP1 lipid biosynthesis pathway proteins. (B, C) *Nrf1α*^*–/–*^cells were transfected with siNC (small-interfering RNA negative control) or siNrf2 (to specifically interfere with Nrf2), before being determined by (B) Western blot analysis of the protein levels as described above or (C) real-time qPCR analysis of their mRNA expression levels. The results were calculated as fold changes (mean ± S.D) with significant decreases (*, *P* < 0.05); statistical significances were determined by using MANOVA. (D, E) *Nrf1α*^*–/–*^cells were transfected with siJUN (to specifically interfere with JUN) or siNC and subjected to Western blotting analysis (D) and real-time qPCR analysis (E), as described above. (F) Wild-type *Nrf1/2*^+/+^cells transiently transfected with expression constructs for JUN (2 μg of DNA) alone or plus FOS (JUN+FOS, each 1 μg of DNA) and then subjected to Western blotting analysis. β-actin was used as a loading control.

Further experiments revealed that silencing of Nrf2 by its specific *siNrf2* (to interfere with its mRNA expression in *Nrf1α*^*–/–*^ cells) also led to obvious down-regulation of AP1 (i.e., a heterodimer of JUN and FOS), HO-1 (a direct co-target of Nrf2 and AP1), and those lipogenic enzymes FASN, ACCα and SCD1 (Fig. 3B, *b1* to *b6*). Next, the real-time qPCR analysis showed that knockdown of *Nrf2* by its *siNrf2* caused significantly reduced expression of *JUN, FASN, ACCα* and *SCD1* (Fig. 3C). This indicates a predominant requirement for Nrf2 in regulating AP1 and the key enzymes in lipid synthesis, besides its cognate target genes *HO-1, NQO1, GCLC* and *GCLM* (Fig. 3C). However, SREBP1 appeared to be roughly unaffected by the silencing of Nrf2 (Fig. 3, B & C).

Interestingly, *siJUN* knockdown of AP1 activity in *Nrf1α*^*–/–*^ cells led to marked decreases in the expression of FASN, ACCα and SCD1 at their protein and mRNA levels (Fig. 3, D & E). However, SREBP1 was almost unaltered by silencing of JUN by siJUN (as shown in this study), although it was shown to regulate activation of specific genes (such as LGALS3) in response to cholesterol loading and hence plays an important role in reprogramming lipid metabolism [25]. Rather, it should also be noted that SREBP1 may not have synergistic regulation with triglycerides-related lipogenic enzymes in this process, albeit lipid droplets are composed of neutral lipids (triglycerides and cholesterol) [26]. Further examination of *Nrf1α*^*–/–*^ cells by transfecting expression constructs for JUN alone or plus FOS revealed that such forced expression of AP1 led to up-regulation of SCD1, but FASN was unaffected (Fig. 3F, *c.f. f5* with *f3*). Intriguingly, only a marginal increase of ACCα was caused by overexpression of JUN alone, but conversely, a substantial decrease of this enzyme was dictated by co-expression of JUN and FOS (Fig. 3F, *f4*). Such dual opposite regulation of ACCα by JUN alone or plus FOS may be determined by a putative functional heterodimer of JUN with another bZIP partner competing together with the AP1 factor, depending on distinct experimental settings.

### 3.4 Nrf1α^−/−^ cell-autonomous induction of the PI3K-AKT-mTOR signaling leads to abnormal hyper-expression of CD36

An important nutritional metabolism mechanism sets into action by the signaling of the PI3K (phosphatidylinositol 3-kinase) to its downstream AKT (also called protein kinase B) and mTOR (mechanistic target of rapamycin kinase) [27]. Upon activation of mTOR by stimulation of nutritional signals, it enables for up-regulation of CD36 (as a key regulator of fatty acid uptake and its transport [28]). As anticipated, a significant increase in the basal constitutive expression of CD36 was found in *Nrf1α*^*–/–*^ cells, when compared with wild-type controls (Fig. 4A, a6). Such aberrant hyper-expression of CD36 was accompanied by dominant cell-autonomous activation of PI3K^P110Δ^ and AKT, as well as of phosphorylated AKT and mTOR at distinct sites (Fig. 4A, a1 to a5). In addition, the loss of *Nrf1α* in *Nrf1α*^*–/–*^ cells led to the basally-augmented expression of a general inflammatory marker COX2 (cyclooxygenase 2, as shown in Fig. 4A, a7); this key enzyme, also called prostaglandin-G/H synthase (PTGS2), is involved in arachidonic acid metabolism. Conversely, Nrf1α-Restored cells gave rise to significant suppression of COX2, CD36, PI3K^p110Δ^, phosphorylated AKT and mTOR, when compared with their counterparts in *Nrf1α*^*–/–*^ cells (Fig. S2, D & E).

**Figure 4.**
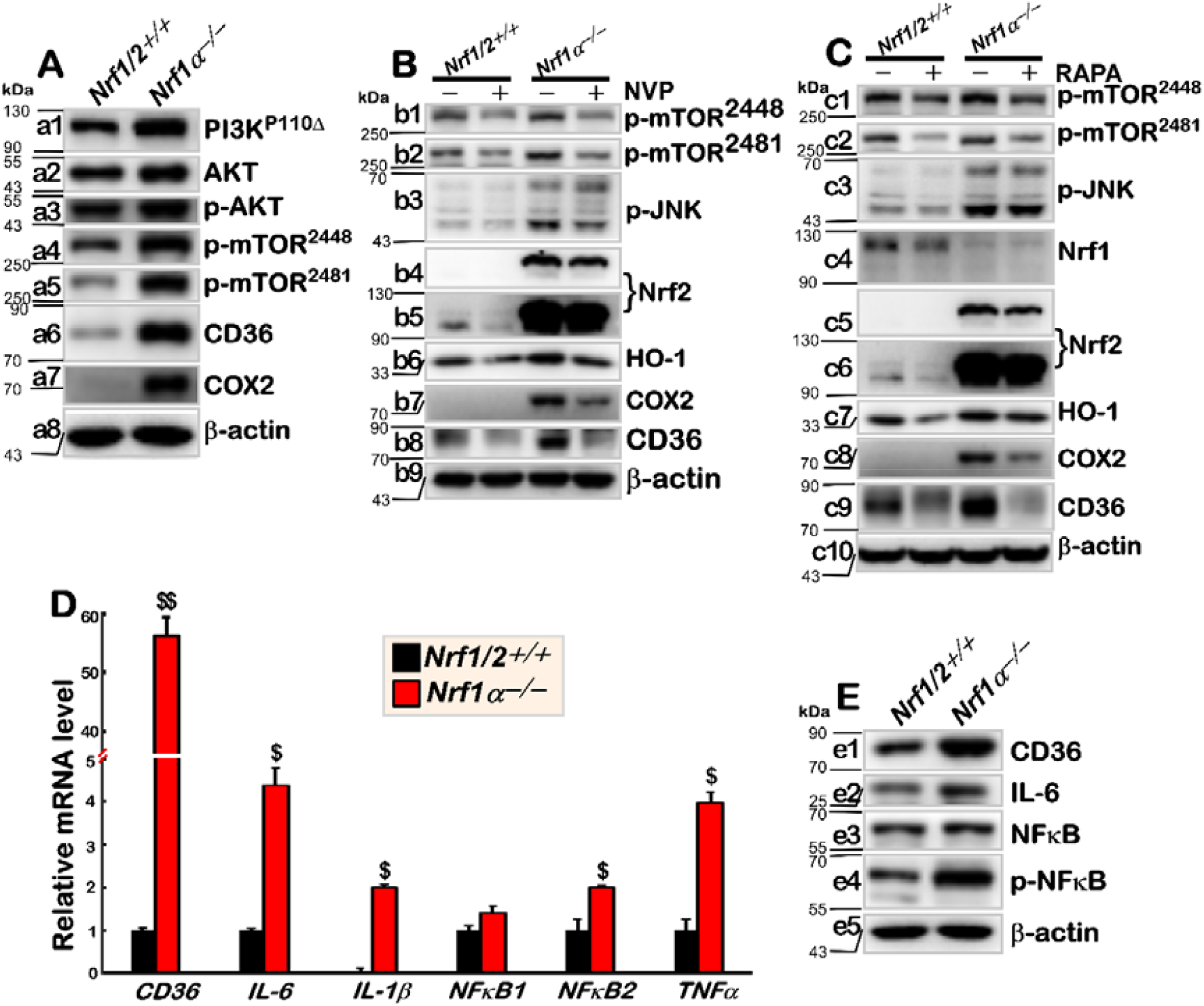
*Nrf1α*^*–/–*^ cell-autonomous up-regulation of CD36 and COX2 through the PI3K-AKT-mTOR sig-naling pathway. (A) Western blotting analysis of indicated protein levels in *Nrf1/2*^+/+^and *Nrf1α*^*–/–*^cell lines as described in the legend for Figure 3(B and C). (B) *Nrf1/2*^+/+^and *Nrf1α*^*–/–*^cells were treated with 0.5 μmol/L of the PI3K-specific inhibitor NVP-BKM120 or (C) 200 nmol/L of the mTOR inhibitor rapamycin (RAPA) for 24 h before indicated protein levels were determined by Western blotting analysis. (D) The mRNA expression levels of those key inflammatory factors and cytokines in *Nrf1/2*^+/+^and *Nrf1α*^*–/–*^ cell lines were measured by real-time qPCR analysis. The results were calculated as fold changes (mean ± S.D) with significant increases ($, *P* < 0.01; $$, *P* < 0.001). The statistical significances were determined by using the Student’s *t*-test or MANOVA. (E) Western blotting analysis of inflammatory factors and cytokines revealed changes in their protein expression levels in *Nrf1/2*^+/+^and *Nrf1α*^*–/–*^cell lines. β-actin was used as a loading control.

To determine whether *Nrf1α*^*–/–*^ cell-autonomous activation of the PI3K-AKT-mTOR signaling is required for up-regulation of CD36 by its deficiency, a PI3K-specific inhibitor NVP-BKM120 (Fig. 4B) and another mTOR-specific inhibitor rapamycin (RAPA, Fig. 4C) were used for treatment of *Nrf1α*^*–/–*^ and *Nrf1/2*^+/+^ cell lines, respectively. As expected, the results revealed that two phosphorylated protein levels of mTOR were substantially diminished by NVP-BKM120 and RAPA in their treated cell lines (Figs. 4B, *b1* & *b2* and *4C, c1* & *c2*). The basal expression levels of COX2 and CD36 were also almost abrogated by NVP-BKM120 and RAPA (Figs. 4B, *b7* & *b8* and 4C, *c8* & *c9*). However, no or less effects of both inhibitors on all other examined proteins (e.g., JNK, Nrf1, Nrf2 and HO-1) were determined (Figs. 4B, *b3* to *b6* and 4C, *c3* to *c7*). This demonstrates that loss of Nrf1α enables to cause a non-controllable activation status of CD36, as well as COX2, primarily by the PI3K-AKT-mTOR signaling pathway.

Since such up-regulation of CD36 can also assist the Toll-like receptor 4 (TLR4) with synergistic activation of the inflammatory factor NFκB and cytokines IL-6, IL-1β and TNFα [29], next we determine whether their changed expression levels in *Nrf1α*^*–/–*^ cells have relevance to the hyper-expression of CD36. The results unveiled that the dramatic activation of CD36 by loss of Nrf1α was accompanied by varying enhancements of NFκB, IL-6, IL-1ß, and TNFα at their protein and/or mRNA expression levels in *Nrf1α*^*–/–*^ cells, when compared with those equivalents of *Nrf1/2*^+/+^ cells (Fig. 4, D & E). In addition, the recovery of Nrf1α in *Nrf1α*^*–/–*^ cells led to apparent decreases in the expression of NFκB and IL-6 (Fig. S2E). Overall, it is postulated that *Nrf1α*^*–/–*^-disordered lipid metabolism may be attributable to aberrant activation of CD36, along with a concomitant inflammatory response.

### 3.5 The inflammatory accumulation of lipids and ROS in Nrf1α^−/−^ cells is alleviated by 2-bromopalmitate

As shown schematically in Fig. 5A, 2BP is a palmitic acid analogue with one bromine (Br) atom at the β-C position, allowing for dramatic improvement of its stability that is more than that of the prototypic palmitate. The physicochemical property of 2BP enables this palmitic acid analogue to block biological processes in which palmitic acids should have participated. For example, 2BP was employed as a natural ligand of PPARδ receptor in fatty acid oxidation [30]. Here we found that triglyceride (TG) accumulation in *Nrf1α*^*–/–*^ cells was significantly inhibited by 2BP (Fig. 5B). Similar treatments with 2BP caused almost abolishment of basal CD36 expression in both *Nrf1α*^*–/–*^ and *Nrf1/2*^+/+^ cell lines (Fig. 5C), and also enabled for substantial suppression of lipid droplets in *Nrf1α*^*–/–*^ cells to considerably lower levels (Fig. 5D). These findings demonstrate that 2BP can alleviate severe lipid deposition mediated by CD36 in *Nrf1α*^*–/–*^ cells.

**Figure 5.**
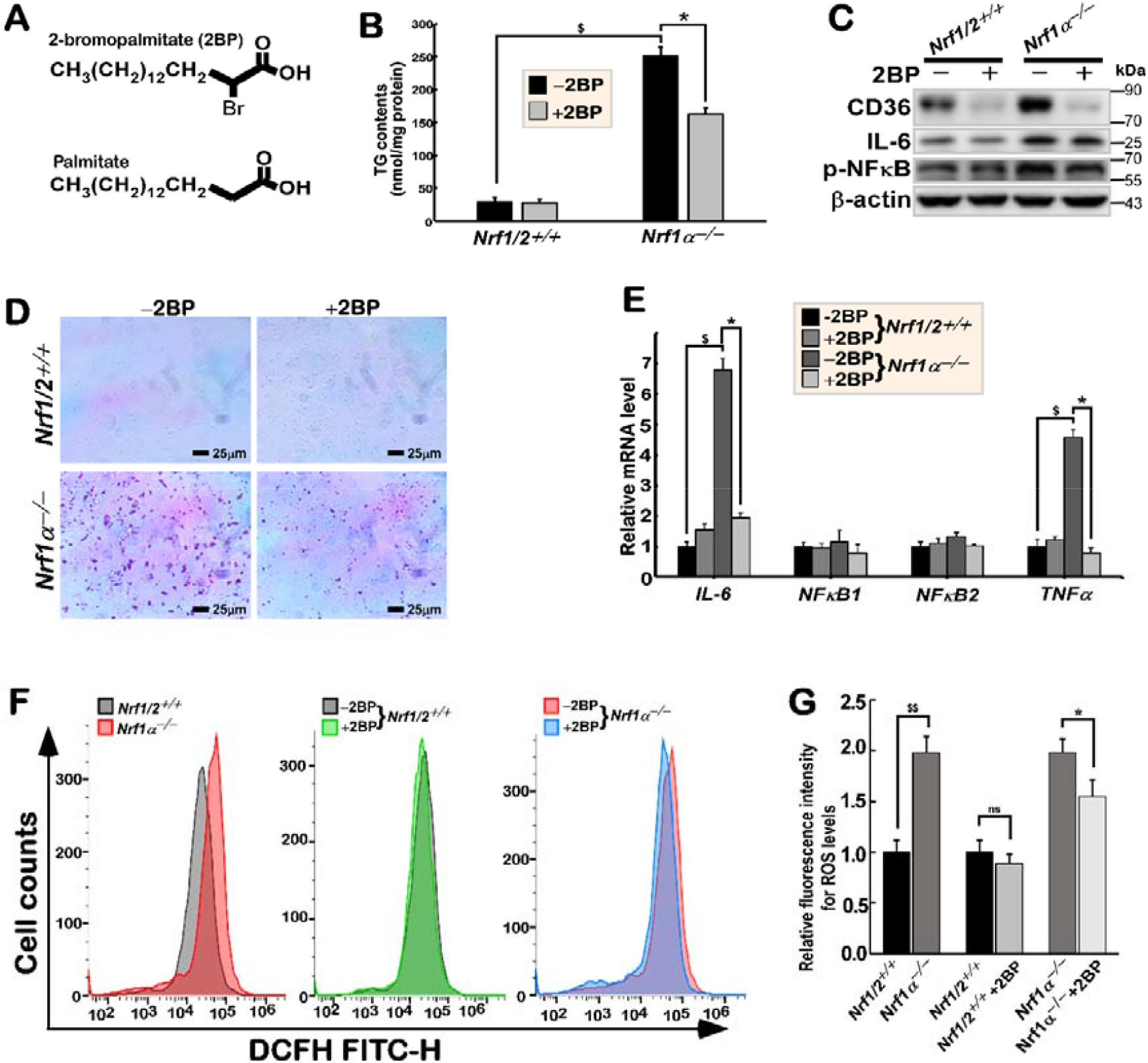
Accumulation of lipids and ROS along with the inflammatory response in *Nrf1α*^*–/–*^ cells are substantially alleviated by 2BP. (A) The schematic shows the molecular structures of palmitic acid (PA) and 2-bromopalmitate (2BP). (B) The triglyceride contents were measured following treatment of *Nrf1/2*^+/+^ and *Nrf1α*^*–/–*^cell lines with 50 μmol/L 2BP. (C) 2BP-treated cell lines were subjected to Western blotting analysis of CD36, IL-6 and p-NFκB. (D) Oil red O staining of *Nrf1/2*^+/+^and *Nrf1α*^*–/–*^cell lines that had been treated with or without 2BP (scale bar = 10 μm). (E) Real-time qPCR analysis of *IL*-6, *NFκB1, NFκB2* and *TNFα* mRNA expression levels. The data were calculated as fold changes (mean ± S.D) with a significant increase ($, *P* < 0.01, relative to the wild-type controls), and another significant decrease (*, P < 0.05, when compared to those measured from the untreated cells). (F) Flow cytometry determination of ROS in *Nrf1/2*^+/+^and *Nrf1α*^*–/–*^cell lines that had been treated with 2BP or without this this palmitic acid analogue, as described in the section of “Materials and Methods”. the results were also graphically shown in (E), along with statistical significances determined by using MANOVA (*, *P* < 0.05; $$, *P* < 0.001; ns, no significance).

Further examination revealed that transcriptional expression levels of the inflammatory cytokines IL-6 and TNFα in *Nrf1α*^*–/–*^ cells were significantly suppressed or even completely abolished by 2BP when compared to wild-type controls (Fig. 5E). Also, the basal IL-6 abundance and active phosphorylated NFκB were repressed by 2BP (Fig. 5C). This implies that the inflammatory response to the loss of *Nrf1α*^*–/–*^ may be prevented by 2BP. Flow cytometry analysis showed a marked accumulation of the intracellular ROS (involved in the inflammatory response) in *Nrf1α*^*–/–*^ cells (Fig. 5F, left panel, with an enhanced peak shifted to the right), but such severe ROS accumulation was also obviously mitigated by 2BP (Fig. 5F, right panel, with a reduced peak shifted to the left). However, no similar ROS changes were observed in *Nrf1/2*^+/+^ cells (Fig. 5F, middle panel). Statistical analysis of these data are shown graphically in Fig. 5G. However, ROS accumulation in the mitochondria of *Nrf1α*^*–/–*^ cells (measured using a MitoTracker Red CMXRos ROS test kit) was only partially reduced by 2BP (Fig. S2F). Collectively, these indicate that 2BP can enable effective alleviation of the inflammatory accumulation of lipids by enhanced biosynthesis and uptake of fatty acids through up-regulation of CD36 in *Nrf1α*^*–/–*^ cells. This occurs concomitantly with a reduction of ROS by 2BP to mitigate the inflammatory response caused by loss of Nrf1α.

### 3.6 The expression of key genes governing lipid catabolism is transcriptionally regulated by Nrf1 through the ARE sites

Clearly, the consensus *ARE* sequence (5′-TGAC/GnnnGC-3′) is known as a *cis*-acting enhancer that drives transcription of genes regulated by Nrf1 and Nrf2 in response to changes in the cellular redox status (i.e., increased production of free radical species including ROS from disordered cell metabolism) or to pro-oxidant xenobiotics that are thiol reactive and mimic oxidative insults [31]. Nrf2 was well documented as an interface between redox and intermediary metabolism [32], so as to protect the liver against steatosis by inhibiting lipogenesis and promoting fatty acid oxidation, but also to exerts its dual role in nonalcoholic fatty liver disease [33]. This may be explained by activation of *ARE*-driven transcription factors that are responsible for regulating the adipocyte differentiation and adipogenesis (e.g., CCAAT/enhancer-binding protein β, peroxisome proliferator-activated receptor-γ, aryl hydrocarbon receptor, and retinoid X receptor-α) and by protection against redox-dependent inactivation of metabolic enzymes (e.g., 3-hydroxy-3-methylglutaryl-CoA reductase). Besides, Nrf1 was also shown to mediate the expression of lipid metabolism-related genes *Lipin1, PGC-1β* [34] and *CD36* [35] by directly binding to the consensus *ARE* sites in their regulatory regions. Here, ChIP-sequencing date obtained from the Encode database (Fig. 6E) showed several peak signals representing the activity of Nrf1 binding to multiple *ARE* sites located within the promoter regions of *ATGL, PGC-1α, PPARα* and *PPARγ*. Such Nrf1-binding details of the ChIP-sequecing data were also provided (Fig. S3, A to D). Further experimental results demonstrated that co-expression of Nrf1 may lead to a certain transactivation activity of some *ARE*-driven luciferase reporter genes (Fig. 6, B to E). These ARE sequences were selected from within the 4k-bp promoter regions of *ATGL, PGC-1α, PPARα* and *PPARγ*, each of which was subcloned into the pGL3-Promoter vector to give rise to those indicated *ARE*-Luc reporter genes as indicated (in Fig. 6F and Table 2). However, Nrf1-mediated transcriptional ability of *ARE*-Luc reporter genes was modestly augmented to less than two-fold changed degrees. This implies that transcriptional expression of lipid catabolism-related genes may be finely tuned by Nrf1 within a certain robust homeostatic extent.

**Figure 6.**
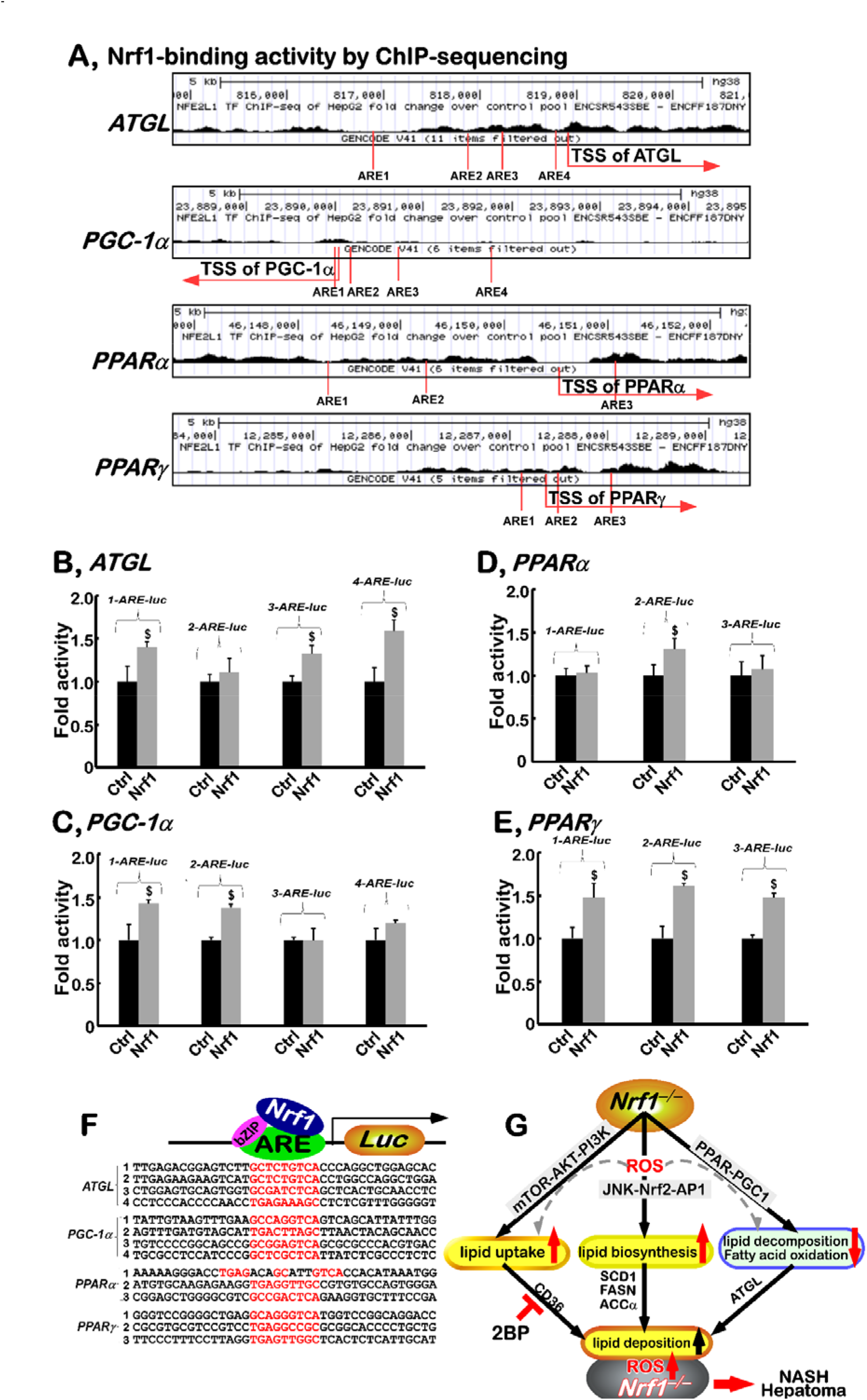
Nrf1 mediates activation of ARE-driven luciferase reporters, along with a proposed model to explain the molecular mechanisms underlying the inflammatory accumulation of lipids and ROS. (A) The ChIP-sequencing date of Nrf1 (i.e., Nfe2l1, obtained from the Encode database) in the promoter regions of *ATGL, PGC-1α, PPARα* and *PPARγ* (the details of which were also deciphered in supplemental Figure S3). The transcription starting sites and multiple *ARE* sites in the gene promoter regions are marked. (B to E) *Nrf1/2*^+/+^ HepaG2 cells were co-transfected with each of those indicated *ARE*-*Luc* or non-*ARE*-*Luc* (used as a background control) reporter plasmids, together with an expression construct for Nrf1 or an empty pcDNA3 vector (serves as an internal control). Then luciferase activity was normalized to their internal p-*RT* reporter controls and also corresponding backgrounds obtained from co-transfection with non-*ARE* reporter and each of the indicated expression constructs. The data were calculated as a fold change relative to the basal activity (at a given value of 1.0) obtained from the transfection of cells with the empty pcDNA3 control and each *ARE*-driven luciferase plasmid. All the data shown herein are representative of at least three independent experiments undertaken on separate occasions that were each performed in triplicates. Significant differences in *ARE*-driven transactivation by Nrf1 were subjected to statistical analysis by using the Student’s *t*-test or MANOVA; significant increases ($, p < 0.01) were shown, by comparison to their basal levels. (F) Fourteen of the indicated ARE-adjoining sequences from lipid catabolism related gene promoters were inserted into the pGL3-Promoter vector to give rise to ARE-Luc reporter genes. (G) A model was proposed for the molecular mechanisms underlying the inflammatory accumulation of lipids and ROS in *Nrf1α*^*–/–*^cells. This schematic provides an overview of the molecular mechanisms by which the inflammatory accumulation of lipids and ROS occurs in *Nrf1α*^*–/–*^cells. Notably, *Nrf1α*^*–/–*^cell-autonomous induction of the PI3K-AKT-mTOR signaling to up-regulation of CD36 results in augmented uptake of fatty acids. The de novo biosynthesis of lipids is up-regulated by the JNK-Nrf2-AP1 signaling to transactivation of their target genes encoding those key metabolic rate-limiting enzymes and their transcription factors, while the lipid oxidative decomposition pathway is down-regulated by the PPAR-PGC1 signaling cascades. Altogether, these result in the inflammatory accumulation of lipids and ROS in *Nrf1α*^*–/–*^cells, but this pathological process is alleviated substantially by 2-bromopalmitate (2BP) to down-regulate the expression of CD36 regulated by Nrf1α.

## 4. Discussion

In the present study, we have provided new evidence unraveling the different or even opposite effects of Nrf1 and Nrf2 on the human lipid metabolism, in addition to our previously reported effects on glucose metabolism[19, 36], the protein homeostasis (proteostasis) [17, 37] and the redox balance and signaling control [38-40]. Thereby, it is of crucial significance to note that the experimental cell model of *Nrf1α*^*–/–*^ with pathophysiological phenotypes appears to largely resemble those manifested by the liver-specific *Nrf1* knockout mice. Such loss of *Nrf1*^*–/–*^ results in the spontaneous development of NASH and ultimately liver cancer, which is attributed to the rapid accumulation of a large volume of lipids, as a key etiology, in the pathological process [9]. Herein, our experimental evidence demonstrated that lipids (i.e., triglyceride contents) were markedly deposited in distinct sizes of the intracellular lipid droplets and widely distributed within the human *Nrf1α*^*–/–*^ hepatoma cells, particularly following treatment with oleic and palmitic acids (Fig. 1, and also see Fig. S4). By contrast, there was a reduced abundance of lipid droplets in *Nrf2*^*–/–*^ cells, even after they were treated with oleic acids.

Interestingly, we have also discovered that abnormal lipid deposition in *Nrf1α*^*–/–*^ cells was substantially blocked by a palmitic acid analogue, i.e., 2-bromopalmitate (2BP, Fig. 5), through significant down-regulation of CD36 (acting as an essential translocator of fatty acids). The concomitant inflammatory response and oxidative stress were ameliorated significantly by 2-bromopalmitate. As a preventive consequence, those key inflammatory cytokines and the transcription factor NFκB were markedly down-regulated, following reduction of ROS and lipids, in 2BP-treated *Nrf1α*^*–/–*^ cells. However, it should also be noted that in addition to the secondary production of ROS stimulated by the accumulation of lipids, the primary production of ROS was partially caused by *Nrf1α*^*–/–*^ cell-autonomous reprogramming of the redox metabolism [19, 36, 39]. Thereby, such portion of *Nrf1α*^*–/–*^ cell-autonomous yield of ROS (particularly in the mitochondria) is inferable to be largely unaffected by 2BP (Figs. 5F & S2F). The evidence presented herein also shows that, even though the putative concomitant ROS resulting from severe lipid deposition may be diminished by 2BP (as an irreversible inhibitor of many membrane-associated enzymes), the oxidative stress is not completely mitigated and recovered to the levels measured from wild-type cells. This palmitic acid analogue 2BP was originally identified as an inhibitor of β-oxidation, but further studies reported that it can also inhibit fatty acyl CoA ligase, glycerol-3-phosphate acyltransferase, mono-, di- and tri-acylglycerol transferases [41]. Therefore, 2BP is thought to block protein palmitoylation by inhibiting such a family of those conserved protein acyl transferases (PATs). Notably, 2BP prevents the incorporation of palmitate into the putative palmitoylated proteins, which may actually reflect alkylation by 2BP at the palmitoylation site, but also seems to reduce palmitoylation of indicated proteins. When used to describe an alkynyl analogue, 2BP also serves a non-selective probe with other targets beyond the palmitoyl transferases [42]. Given the ubiquitous usage of 2BP in a large number of studies published, it should have a significant ramification on the palmitoylation area [43]. For instance, CD36 palmitoylation causes an increase in its trafficking and also cell-surface location of adipocytes, which makes oleate uptake easier [44]. Once palmitoylation of CD36 is inhibited, its hydrophobicity and accessibility to the plasma membrane are accordingly decreased, which results in less binding and absorption of long-chain fat acids [45]. This implies that palmitoylated CD36, as a cognate key target of Nrf1, plays a crucial role in fat acid and other lipid metabolism. Consistently, 2-BP was de *facto* employed to effectively prevent CD36 palmitoylation modification [46].

Here, we have also provided evidence revealing that abnormal accumulation of lipids in *Nrf1α*^*–/–*^ cells resulted from incremented uptake of fatty acids through constitutive activation of the PI3K-AKT-mTOR signaling pathway leading to up-regulation of CD36. The CD36-augmented uptake of lipids is fully consistent with the results obtained from the tissue-specific *Nrf1*^*–/–*^ mice [10, 35]. These two independent groups further showed that Nrf1 is essential for controlling the translocation of fatty acids and the cystine/cysteine contents by governing a complex hierarchical regulatory network in redox and lipid metabolism. In addition to CD36, there are also some members of the cytochrome family, such as Cyp 4A, were also up-regulated by the loss of *Nrf1*^*–/–*^, thus leading to the generation of ROS from the oxidative metabolism of fatty acids, in the murine deficient liver cells [10, 11]. Besides, we also previously showed that Nrf1 and Nrf2 contribute to dual opposing regulation of the PTEN-AKT signaling to be selectively repressed or activated in the pathogenesis of NASH and liver cancer [17]. In the present study, we further validated that the extended PI3K-AKT-mTOR signaling to the inflammatory marker COX2, as well as the key inflammatory responsive factors along with CD36, was also constitutively activated by such loss of Nrf1α in human hepatoma cells.

In this study, we have provided more insight into the molecular mechanisms of how *Nrf1α*^*–/–*^ deficiency is involved in lipid metabolism disorders, along with the relevant pathway of enzymatic reactions leading to lipid deposition during development of NASH and hepatoma. Our evidence demonstrates that the key enzymes FASN, SCD1 and ACCα and the transcription factor SREBP1 (involved in the de novo biosynthesis of lipids) were significantly up-regulated in *Nrf1α*^*–/–*^ cells (with hyper-active Nrf2 retained). However, they were unaffected or even down-regulated by inactivation of *Nrf2 in Nrf2*^*–/–*^ cells. Further experimental evidence showed that such up-regulation of the lipid biosynthesis pathway by *Nrf1α*^*–/–*^ deficiency may occur through the JNK-Nrf2-AP1 signaling. An explanation for the results regarding Nrf2 may be attributed to the heterodimeric formation of Nrf2 with each of sMaf or other bZIP proteins, thereby competing with JUN and FOS against another cognate heterodimeric AP1 factor for selective physical binding to the potential ARE consensus sites within their target gene promoter regions [47].

We also found that ATGL and the nuclear receptor PPAR-PGC1 signaling towards the lipid oxidative decomposition pathway were markedly down-regulated in *Nrf1α*^*–/–*^ cells, but they were unaffected or even up-regulated in *Nrf2*^*–/–*^ cells. The activation of the PPAR-PGC1 signaling was also supported by the evidence obtained from the tissue-specific *Nrf1*^*–/–*^ mice [12, 48]. Hirotsu et al [12] showed that liver-specific *Nrf1* deficiency cannot only lead to decreased expression of Lipin1 and PGC-1β, but conversely, the presence of Nrf1 enabled the CNC-bZIP factor to directly mediate transcriptional activation of Lipin1 and PGC-1β by binding to the ARE sequences in their promoter regions. Collectively, this indicates that Nrf1 is predominantly essential for regulating the critical genes involved in lipid catabolism. Upon loss of *Nrf1α*^*–/–*^, constitutive activation of the PPAR-PGC1 signaling cascades is implicated in the severe aberrant accumulation of lipids. The resulting metabolic pathological process may be significantly prevented by 2BP, because this palmitic acid analogue has also been shown to serve as a ligand of PPARδ receptor involved in the fatty acid oxidation [30], although the detailed molecular mechanisms remain unclear.

In summary, this work demonstrates that both Nrf1 and Nrf2 make different or even opposite contributions to the intracellular lipid metabolism, as illustrated in a model (Fig. 6G). Upon knockout of Nrf1α, the de novo biosynthesis of lipids is incremented by significant induction of the JNK-Nrf2-AP1 signaling pathway in *Nrf1α*^*–/–*^ cells, whereas the lipid oxidative decomposition pathway was down-regulated by the PPAR-PGC1 signaling cascades. *Nrf1α*^*–/–*^ cell-autonomous activation of the PI3K-AKT-mTOR signaling towards up-regulation of CD36 leads to augmented uptake of fatty acids. Taken altogether, these results lead to severe accumulation of lipids deposited as lipid droplets. By contrast, *Nrf2*^*–/–*^ cells gave rise to rather decreases in both lipid synthesis and uptake capacity, along with enhanced oxidative metabolism of fatty acids. Importantly, such *Nrf1α*^*–/–*^-leading lipid metabolism disorders and the relevant inflammatory responses are substantially rescued by 2BP, probably through blocking palmitoylation of CD36 (as a target of Nrf1). Overall, this study provides a novel potential strategy for chemoprevention of the inflammatory transformation into malignant cancer and the relevant treatment by precision targeting of Nrf1, Nrf2 alone or both in translational medicine.

## Author contributions

R.D. repeated most of the experiments that Z.Z. originally performed together with M.W., S.H. and J.F., collected the data with their original ones being edited, wrote the draft of this manuscript with most figures and supplemental information, and also revised this manuscript. Both P.M. and Z(W).Z. helped to wrote and discuss the manuscript. Lastly, Y.Z. designed and supervised the study, analyzed all the data, helped to prepare all figures and cartoons, wrote and revised the paper.

## Acknowledgments

We are greatly thankful to Drs. Lu Qiu (Zhengzhou University, Henan, China) and Yonggang Ren (North Sichuan Medical College, Sichuan, China) for their involvement in establishing those knockout cell lines used in this study. We are also thankful to all the present and past members of Prof. Zhang’s laboratory (at Chongqing University, China) for giving critical discussion and invaluable help with this work. This study was funded by the National Natural Science Foundation of China (NSFC, with two project grants 81872336 and 82073079) awarded to Prof. Yiguo Zhang.

## Supplemental Figure legends

**Figure S1.**
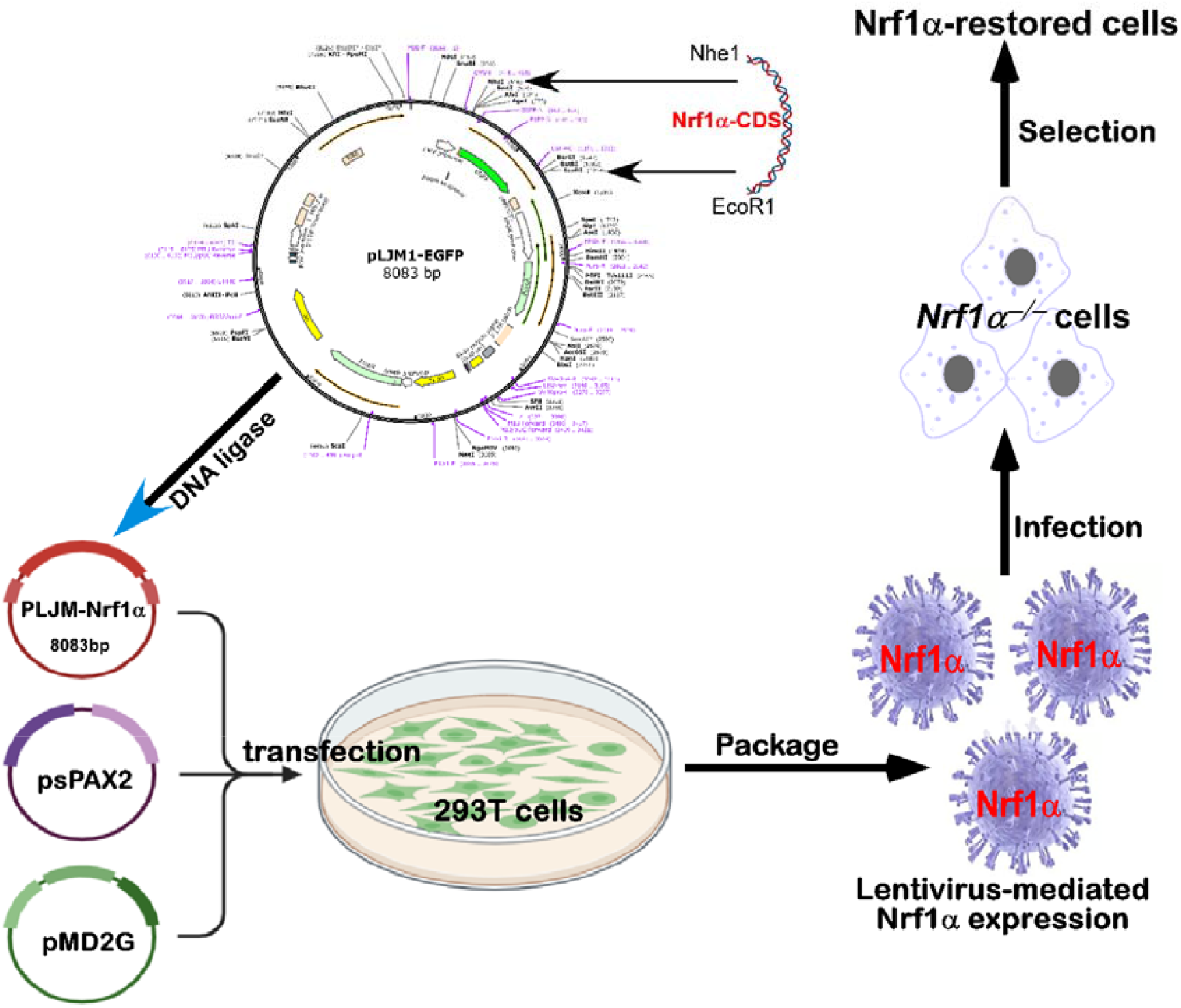
A flow chart for a lentivirus-packaged pLJM1-Nrf1α plasmid that was constructed and then infected into HepG2 cells, before a monoclonal Nrf1α-restored expression cell line was established to allow for stable expression of ectopic Nrf1α.

**Figure S2.**
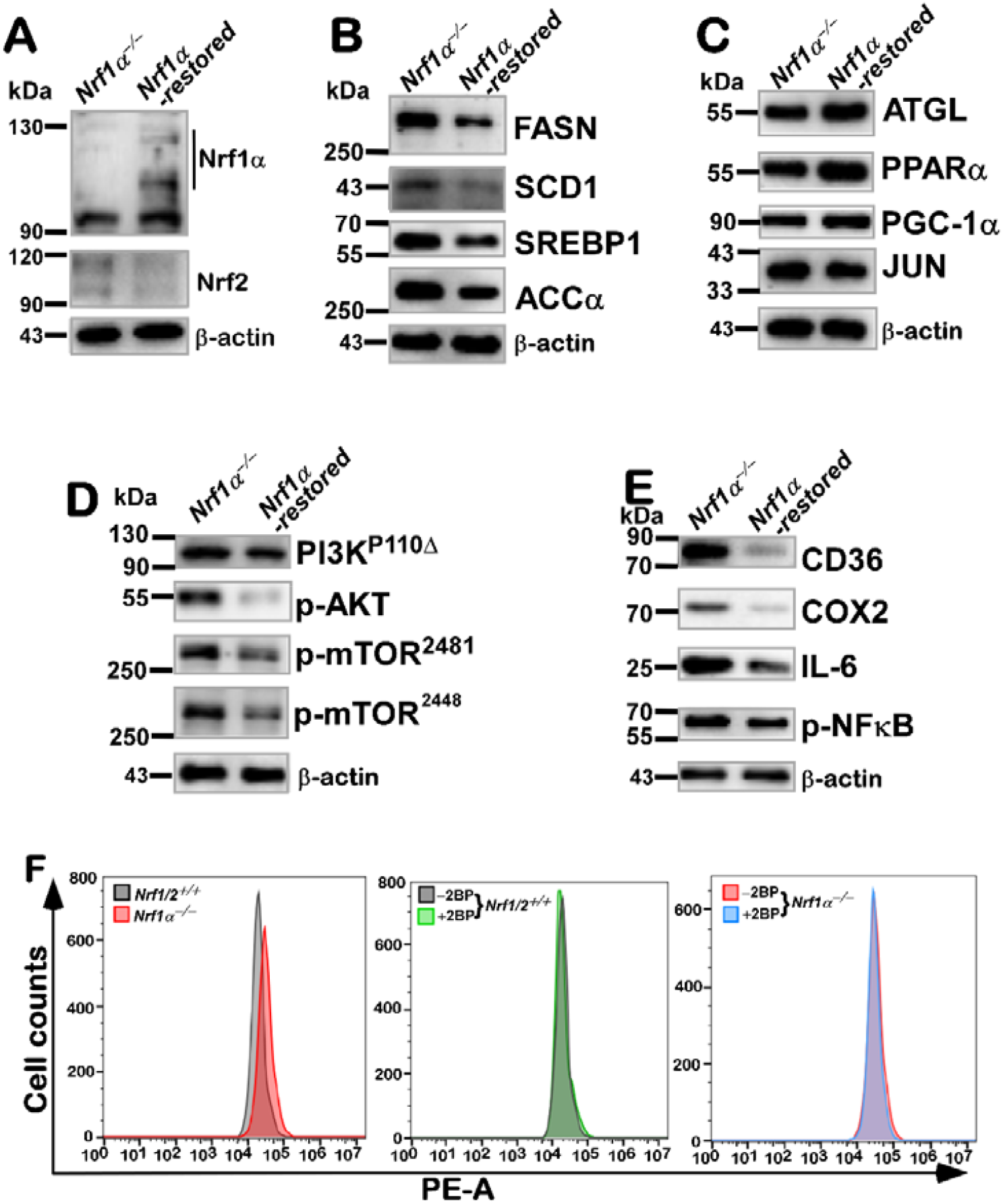
A series of repeatable experiments using the Nrf1α-restored expression cell line, in order to confirm that those changes observed in *Nrf1α*^*–/–*^cells could be rescued by restoring stable expression of Nrf1, so that accumulation of lipids and ROS, along with the inflammatory response in *Nrf1α*^*–/–*^cells are substantially alleviated by 2BP. The expression levels of those examined proteins in the lentivirally-mediated recombinant Nrf1^α^-Restored cell lines were detected by Western blotting with indicated antibodies (A to E); Flow cytometry determination of ROS in *Nrf1/2*^+/+^and *Nrf1α*^*–/–*^cell lines that had been treated with 2BP or without this this palmitic acid analogue, as described in the section of “Materials and Methods”.

**Figure S3.**
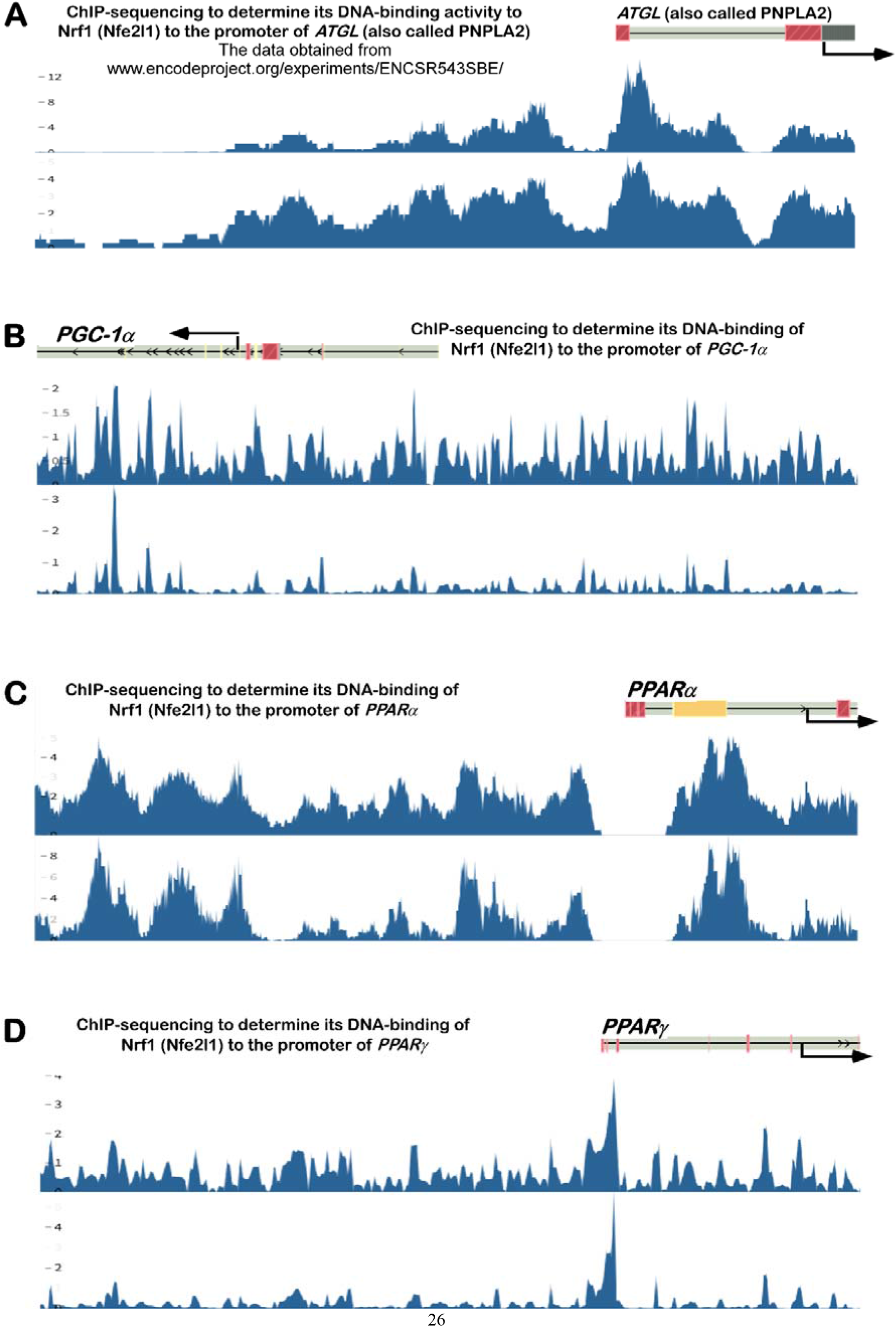
The ChIP-sequencing date of Nrf1 (i.e., Nfe2l1, obtained from the Encode database) binding to the promoter regions of ATGL, *PGC-1α, PPARα* and *PPARγ*.

**Figure S4.**
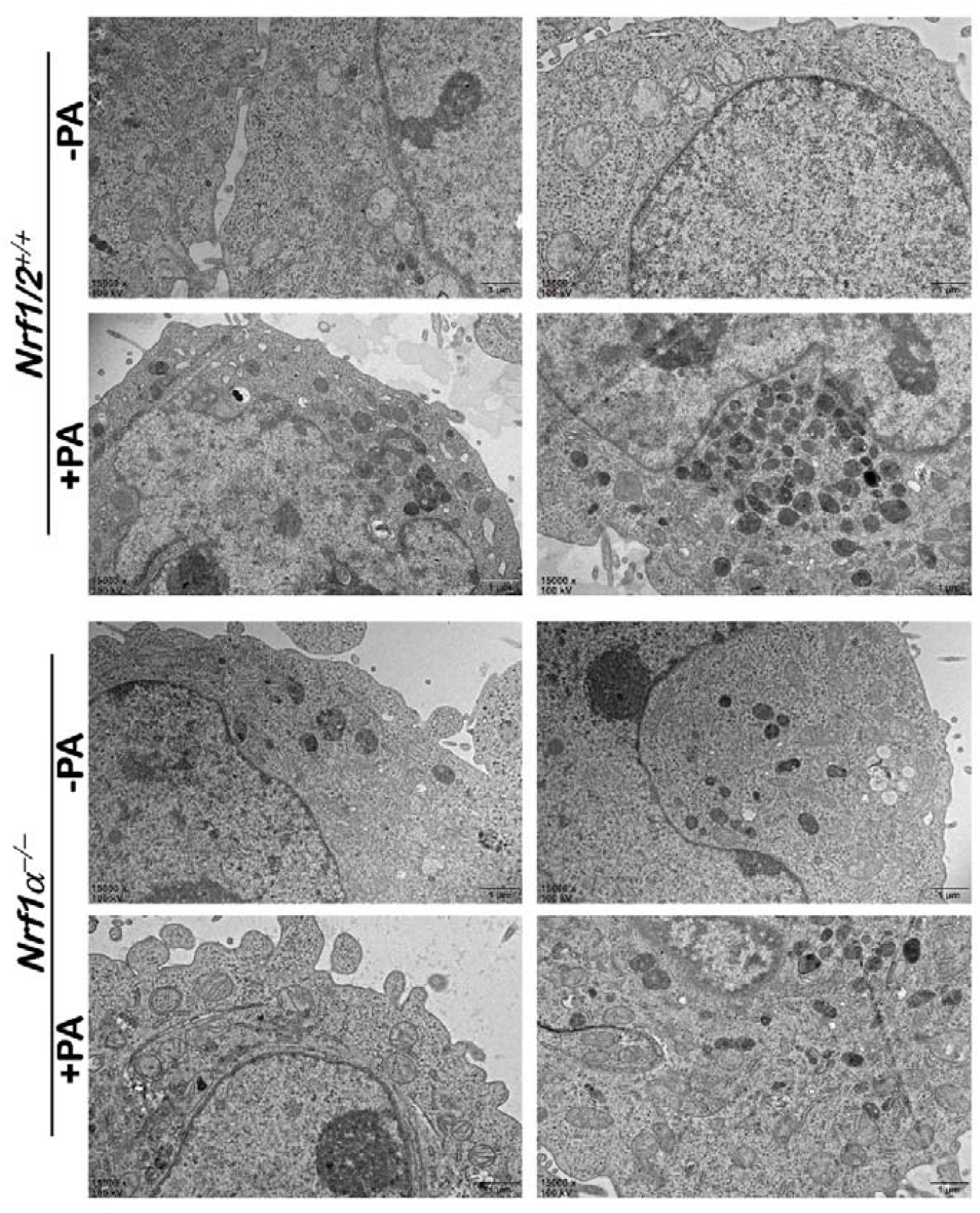
The electronic micrographs representing WT (*Nrf1/2*^+/+^) and *Nrf1α*^*–/–*^cell line that had been treated with palmitic acids. The scale bar = 2 μm in ×2K pictures, or = 1 μm in ×7K pictures, or = 0.25 μm in ×20K pictures.

